# A new high-resolution three-dimensional retinal implant: *System design and preliminary human results*

**DOI:** 10.1101/2022.09.14.507901

**Authors:** Leonid Yanovitch, Dorit Raz-Prag, Yael Hanein

## Abstract

The NR600 retinal prosthetic device is a unique newly developed implant intended to restore visual perception to people who lost their vision due to retinal degenerative diseases. The miniature implant captures the visual image in place of the damaged photoreceptor cells and creates the electrical stimulation required to activate the preserved retinal cell layers. The NR600 system transduces visual signal into electrical signals that are delivered to the retina via an array of needle shaped electrodes to minimize electrical activation levels and improve stimulation localization. NR600 consists of two components: A miniature implantable chip and eyeglasses set worn by the patient. The eyeglasses deliver power and control the implantable device. In this report, we present the NR600 system design, its optical, electrical, and electro-chemical characteristics, and preliminary results from human subjects.

## 1. Introduction

Over 200 million people worldwide suffer from severe vision impairment due to a variety of diseases, including retinal degenerative diseases [1]. The most prevalent retinal degenerative diseases are retinitis pigmentosa (RP), which is the most common inherited cause of blindness in people between the ages of 20 and 60 [2], [3], [4], dry type age-related macular degeneration (AMD), primarily affecting people aged 60 and older [3], [4], [7], [8] and diabetic retinopathy (DR) which gradually degenerate the retina of unbalanced diabetic patients. These conditions cause progressive damage to the retina resulting in different levels of vision impairment.

Retinitis pigmentosa (RP) is a group of genetic disorders that involve a breakdown and loss of cells in the retina resulting in gradual loss of peripheral vision and eventually complete blindness. Since there are many gene mutations leading to RP, its progression can differ greatly from patient to patient. RP typically appears in childhood showing difficulty to adjust to lighting changes and reduced night vision. As the disease progresses, patient’s visual field gradually narrows down. Some patients retain central vision and a restricted visual field into their 50s, while others experience significant vision loss in early adulthood. Eventually, most individuals with RP will lose most of their sight. No effective treatment is currently available for halting the progression of RP.

AMD typically occurs in older adults and results in loss of vision in the center of the visual field (macula). Individuals with AMD may first notice a blurring of central vision, especially during tasks that require detailed vision. Other common symptoms include distorted vision for which straight lines appear wavy, pigmentary changes and photo-stress. As the disease progresses, blind spots may form within the central visual field and in severe cases can lead to complete loss of vision. Treatment for early dry-AMD is generally nutritional therapy, emphasizing certain vitamin groups. For wet-AMD, the most common treatment to date are anti vascular endothelial growth factor (anti-EVGF) injections which inhibit the formation of new blood vessels behind the retina, preventing further vision loss. Late-stage AMD, specifically in those patients who are classified as geographic atrophy (GA), causes severe vision impairment, with no current available treatment other than supportive treatment as external vision aids.

Several clinical trials of retinal prostheses [5] for RP and late-stage GA were conducted and three of them were regulatory approved (Argus II by Second Sight, Alpha I/AMS by Retina Implant GmbH & IRIS-II by Pixium) but are not presently commercially available. The Argus II by Second Sight [6]–[8] and PRIMA by Pixium demonstrated, in early feasibility studies, compatibility between artificial vision and natural peripheral vision in GA patients. Gene-therapy research promises future hope for RP patients, but currently only the pioneer AAV-based LUXTURNA by Spark Therapeutics (now owned by Roche AG) is available for the treatment of RPE-65-related retinal degenerations. The scarcity of treatments combined with the limited performance of contemporary retinal implants, creates an opportunity for a new generation of retinal implants. Such implants can potentially reach the benchmark of functional vision by utilizing denser and more effective electrode arrays.

Over the last several decades several artificial retina devices were developed [9]–[11], some reaching human trials showing some level of phosphene detection and image recognition [7], [12]–[14]. Despite extensive efforts, device performances were so far modest [15]. In this paper we report on a newly developed implant intended to restore sight to people who lost their vision due to retinal degenerative diseases. The miniature NR600 Implant, captures the visual image (taking the place of the damaged photoreceptor cells) and creates the electrical stimulation required to activate the remaining preserved retinal cells. The NR600 retinal prosthesis has several unique properties distinguishing it from previously reported devices. First, it has a dense needle-like penetrating array, which allows efficient coupling between the electrodes and the neural tissue. It is implanted as an epi-retina device yet functions as an intra-retina device. Consequently, it benefits from a simple implantation procedure along with good electrode-tissue coupling. Moreover, the NR600 implant has no internal power source, instead it relies on photovoltaic elements embedded in the implant. The power delivered to the implant by the eyeglasses supports enough energy for stimulation and for powering an internal ASIC device that controls the implant output parameters allowing important features such as an adaptive dynamic range and stimulation signal tuning per patient.

## 2. NR600 Building Blocks and Design Considerations

NR600 consists of two main components (see Figure 1): A miniature implantable unit and a set of eyeglasses worn by the patient. The implantable unit encapsulates a silicon photodiode imager, a PV cell and two ASIC units which convert the output from the imager into specific neurostimulation signals. The eyeglasses supply both power and communication to the NR600 implant while allowing visible image to pass through with minimal distortion. To supply both power and communication effectively, the glasses use Infra-red (IR) modulated illumination. A unique optical design enables steering the IR radiation into the patient’s eye, while maintaining clear visible field in front.

**Figure 1:**
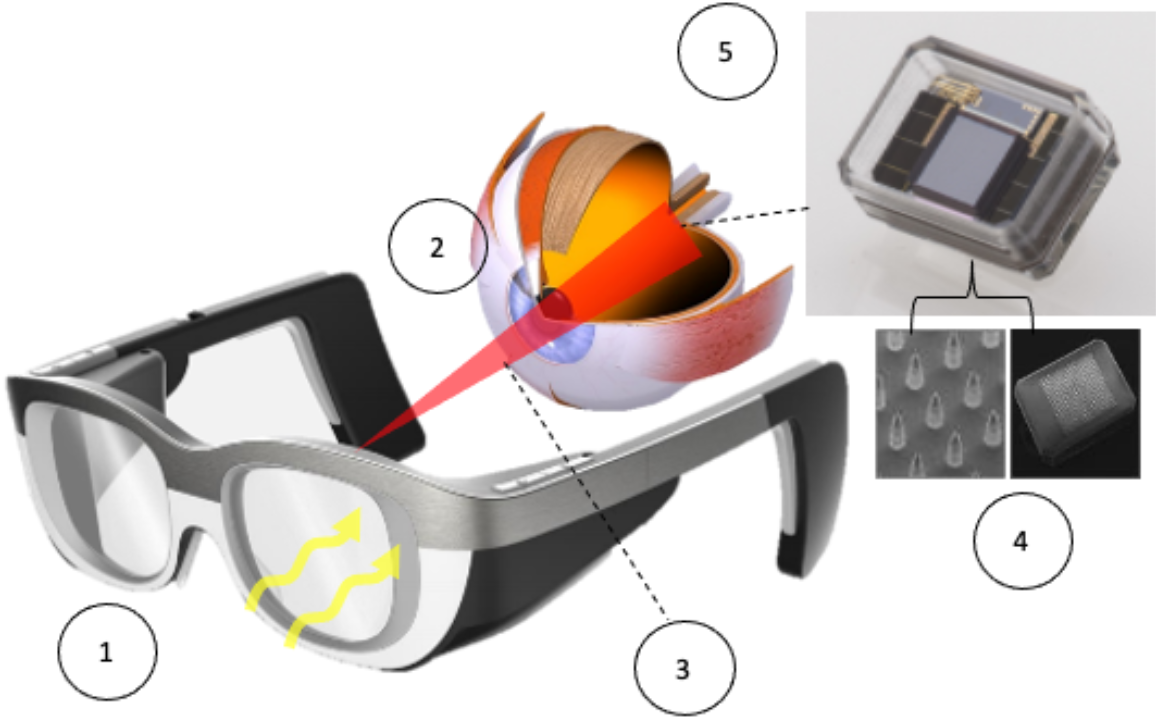
NR600 system: Implant (5) and eyeglasses (1) operate as follows: Visible light passes through the glasses and patient’s IOL. (2) Image is received on the imager located at the retina and light is converted to electrical stimulation. (3) Infra-Red beam powers the implant wirelessly and controls the stimulation parameters (4) Intra-retinal electrodes stimulate retinal neurons to deliver an image to the brain.

### 2.1.1 Electrode Array

Electrical stimulation of NR600 is delivered to the retina using a dense array of intra retinal penetrating microelectrodes. These three-dimensional microelectrodes penetrate the conserved retinal layers that originally received the signal from the photoreceptors and stimulate them electrically. The unique shape and structure of the electrodes enable precise local stimulation of the targeted healthy retinal cells. The array consists of electrically isolated, dielectrically coated, single crystalline silicon (SCS) electrodes. A dielectric barrier between adjacent electrodes runs through the entire substrate and allows excellent electrical isolation and no crosstalk between neighboring electrodes. In order to optimize the electrode-tissue interface for effective stimulation, the electrode tips (i.e. active area) are coated with high specific capacitance TiN coating. The rest of the electrode (i.e. inactive isolated area) is deposited with an SiO_2_/SiC layer which effectively isolates the electrode base from the medium.

### 2.1.2 Structural Properties

The eye curvature and retinal thickness present a challenge for choosing an electrode length that will fit most patients. Nevertheless, imaging tools such as optical coherence tomography (OCT) can be used to screen patients before the implantation and in order to fit them with the best length of electrodes. A recent study on RP patients showed that 150 µm electrodes are on average an optimal length for temporal implantation. In NR600, a typical length of 150±30 µm, and a maximum exposed tip height (TiN, see below) of 50 µm were implemented. The validity of these values for neural stimulation was also validated with electrochemical tests, which will be discussed separately. Typical needle structures are presented in Figure 2.

**Figure 2:**
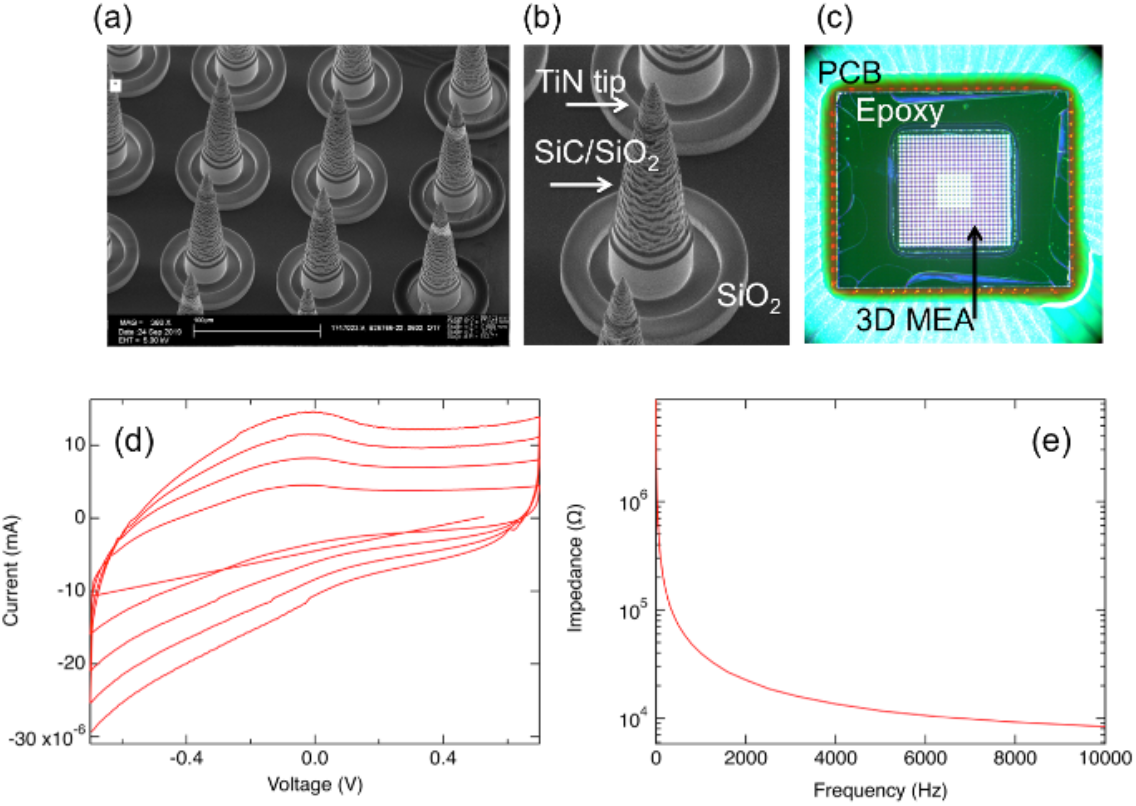
The NR600 3D electrode array. (a, b) SEM images of silicon needles with SiO_2_ trench isolation and exposed TiN tips. (c) Silicon micro electrode array (MEA) mounted on PCB for electrochemical analysis and ex-vivo investigations. (d, e) Typical cyclic voltammetry (CV) and electrochemical impedance spectroscopy (EIS) data of NR600 electrodes. CV scan rates in a were 200, 400, 600 and 800 mV/s.

### 2.1.3 Electrochemical Characteristics

Capacitive charge injection is advantageous over faradic charge injection, as no chemical species are created or consumed during a stimulation phase which ensure high safety of these electrodes for stable neurostimulation. However, as the charge stored in the double layer per unit area is small, high charge injection capacity is possible only with ultra-porous electrodes. Two key examples for purely capacitive materials are porous titanium-nitride (TiN) and doped ultra nano-crystalline diamonds (UNCD).

NR600 uses ultra-porous TiN as the electrode material. Titanium-nitride (TiN) is a chemically stable conductor with good biocompatibility. It is extensively used in cardiac pacing applications and in various neural interfaces. Large charge injection capacity and high charge storage capacity are obtained by fabricating electrodes with high surface roughness, such that the electrochemical surface area (ESA) greatly exceeds the geometrical surface area (GSA). This high ESA to GSA ratio is obtained through the fractal surface area, typically created by reactive sputtering of Titanium (Ti) in Nitrogen (N_2_) and Argon (Ar) gas mixture. The gas mixture ratio, deposition temperature, deposition length, substrate composition, all strongly affect the resulting surface morphology of the TiN coating. Values reported in the literature range from 0.5 to almost 5 mC/cm^2^. Typically, for 1kHz frequency (equivalent to ~1ms pulses), fractal TiN can safely reach a charge injection limit of 2 mC/cm^2^.

The thickness of the applied coating and the grain size have a strong effect on the overall charge storage capacity (CSC). The electrolyte resistance and capacitance on the interior surface of the pore combine to form a transmission-line with a time-constant defined by the pore geometry, electrolyte resistivity and the interfacial double-layer capacitance. As a result of this time-constant the total ESA of the electrode is not accessed. Narrower and deeper pores give rise to higher time-constants, and their charge-injection capacity is more difficult to access than that of electrodes with shallow pores. Increased surface roughness would not guarantee an increase in the charge injection capacity, especially for shorter pulse durations (<1ms). These issues were also considered in the electrode design.

Titanium-nitride (TiN) is a particularly compelling material as it allows both excellent charge injection capability and mechanical/electrochemical long-term stability. For very small electrodes, where the charge injection capacity is of critical importance, activated iridium oxide film (AIROF) and sputtered iridium oxide film (SIROF) coatings will require anodic biasing to maximize their charge injection capability. Long-term stability should also be noted, as imbalanced charge injection may result in Ir ion migration and electrode deterioration.

The electrode electrochemical properties were studied extensively, measuring their voltage transient to current stimuli, their CSC, their specific capacitance and their behavior in electrochemical impedance spectroscopy (EIS) measurements (see Fig. 2). For clarity, we briefly describe these testing techniques. Electrochemical impedance spectroscopy (EIS) involves measuring the electrical impedance and phase angle obtained with sinusoidal voltage or current excitation of the electrode. The measurement is perfromed over a broad frequency range, typically from 1 Hz to 10^5^ Hz, and the magnitude of the excitation is sufficiently small that a linear current-voltage response is obtained, typically ~10 mV.

Using the measured EIS curve, we can obtain the electrical model for the electrode and tissue by means of a simple fitting. Voltage transient technique involves recording the potential on the electrode for a rectangular current pulse. It is sometimes used as an alternative to EIS as it requires very little additional circuitry to implement. Assuming the faradaic leakage of the electrode is very small, the effective capacity of the electrode and the tissue resistance can be extracted from a voltage transient recording.

Charge storage capacity (CSC), the total amount of charge per unit area available from the electrode can be calculated from a Cyclic Voltammetry (CV) measurement by integrating the area underneath the curve. Electrode capacitance can also be extracted from this measurement by dividing the CSC by the potential window. CVs are most often performed at slow sweep rates of few millivolts per second, thus the extracted data is often referred to as DC measurements. At higher sweeping frequencies, both the capacitance and the CSC of an electrode change. Generally speaking, CVs at high sweep rates provide information primarily on exposed areas of the electrode, whereas CVs at slow sweep rates include contributions from more occluded areas of the electrode (e.g. where the passivation overlaps the active area).

### 2.1.4 Electrode Isolation and Passivation Materials

As the electrode array is exposed to body fluids, it is susceptible to corrosion and dissolution. To protect the array and maximize its lifetime, a stable passivation layer is required. This passivation layer is typically deposited along the array surface, as well as on the electrodes shaft, where it helps to minimize shunting currents and limits stimulation to the electrode active area region. Silicon is susceptible to dissolution in a saline environment, with etch rates as high as 67 nm per day or roughly 2 µm per month. Effectively, such dissolution rate will translate to complete array failure within several months. Hence, a passivation layer is required to drastically reduce dissolution rates. A wide variety of passivation materials may be used for chronically implantable arrays, including Polyimide, Parylene, Silicon Oxide (SiO_2_), Silicon Nitride (SiN), Silicon Carbide (SiC), Nano-diamond coatings and more. While all were shown to be suitable for long-term chronic implantation, they still differ by their application methods, electrical properties, water vapor transmission rates and dissolution rates. Finding one material that meets all the requirements for coating neural interfaces is challenging. For example, SiN has relatively high dissolution ratio, while SiC and nano-diamonds need relatively high deposition temperatures. Polyimide and Parylene are easily coated in room temperature but susceptible to cracking and delamination.

Silicon carbide in its crystalline form is a widely used semiconductor, in LEDs, detectors and high temperature integrated circuits. In non-electric applications, amorphous SiC appears as a protective coating against corrosion, moisture, etching and abrasion or for bio-molecular and medical applications. All of these applications are based on unique properties of SiC, such as a wide band gap, high electron mobility, high thermal conductivity, and a high melting point. Amorphous silicon carbide is particularly suited for passivation of chronically implantable devices due to its biocompatibility and negligibly low dissolution rate in saline. NR600 uses a plasma-enhanced chemical vapor deposition (PECVD) SiC coating for passivation needs.

To conclude, test units were characterized using EIS and CV techniques, and electrical properties were extracted using dedicated Matlab script (Figure 2 and Table 1). Eight 18-electrode arrays were characterized. The charge injection capacity and Randle’s model parameters (C_E_ and R_E_) were extracted. Acceleration by increased stimulation frequency was used to validate under overstressed experiments.

**Table 1:** NR600 electrochemical parameters.

| Parameter | Value |  | Acceptance criteria | Comments |
| --- | --- | --- | --- | --- |
| Charge injection capacitance [nC] | 1.63 | $\pm 0.22$ | $>1$ | Ex-vivo experiments using the 3D electrodes showed typical stimulation threshold of 0.5-1 nC |
| Capacitance - $C_E$ [nF] | 3.09 | $\pm 0.40$ | $>2$ | |
| Faradaic resistance - $R_E$ [Mohm] | 17.93 | $\pm 3.38$ | $>10$ | Driver compliance voltage is 0.8 V, so no more than $\sim 0.05 \mu A$ will be lost to faradaic leakage. This is negligible relative to the stimuli amplitude which is above $1 \mu A$. |

## 2.2 NR600 Electronics

The implant consists of several main building blocks depicted in the block diagram in Fig. 3. Primarily, the implant consists of: (1) An imager, which captures the natural image impinging at the back of the eye; (2) A low power processor that translates the image to an electrical stimulation signal; (3) PV cells supplying power to the implant; (4) Receiver (RCVR) for controlling implant configuration parameters, such as pulse duration and amplitude; and (5) A digital low pass filter. These circuits are also designed to guarantee proper power and voltage management, charge balancing and compatibility with a safe electrode water window. Below is a concise summary of some of these elements.

**Figure 3:**
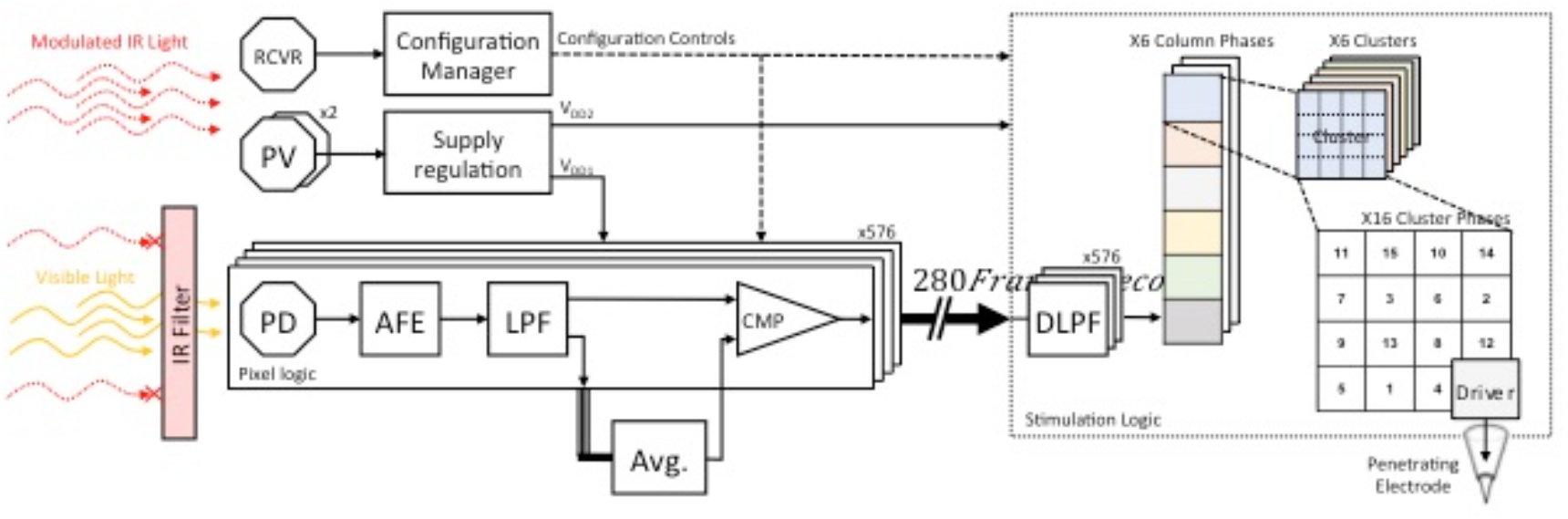
NR600 implant block diagram.

### 2.2.1 Charge Injection and Balancing

The NR600 implant has a proprietary circuitry design that ensures that for any given stimulating electrode, it will be surrounded by multiple return electrodes. This special design provides local return for better localizing the stimulation. In addition, the local return increases the gradient of the generated electromagnetic field during the stimulation phases allowing neural activation at lower charge thresholds. In Figure 5 we plot a schematic representation of the neurostimulator circuitry implemented in the NR600 implant.

The implemented current-mode driver is the most common driver type for neural stimulation, where the charge is delivered by a constant current source with its level easily controlled by the stimulation duration. Voltage-mode drivers are much simpler, as they apply voltage directly on the electrode for a defined duration. Unfortunately, the simplicity comes on the expense of accuracy, as the amount of charge driven depends on electrode impedance and cannot be directly quantified. Finally, charge-mode drivers rely on fast and successive switching of charge from a discrete capacitor to the electrode. Using this method, the driven charge is quantifiable, while retaining higher efficiency compared to current-mode drivers. Nevertheless, charge-mode drivers are not very common as they are generally more complicated and require more silicon area to implement. Additionally, for accuracy across a wider range (1 nC – 100 nC for example), charge-mode drivers require more than one capacitor on which charge is collected. In Fig. 4a schematic representation of the charge injection circuitry implemented in the NR600 implant is plotted.

**Figure 4:**
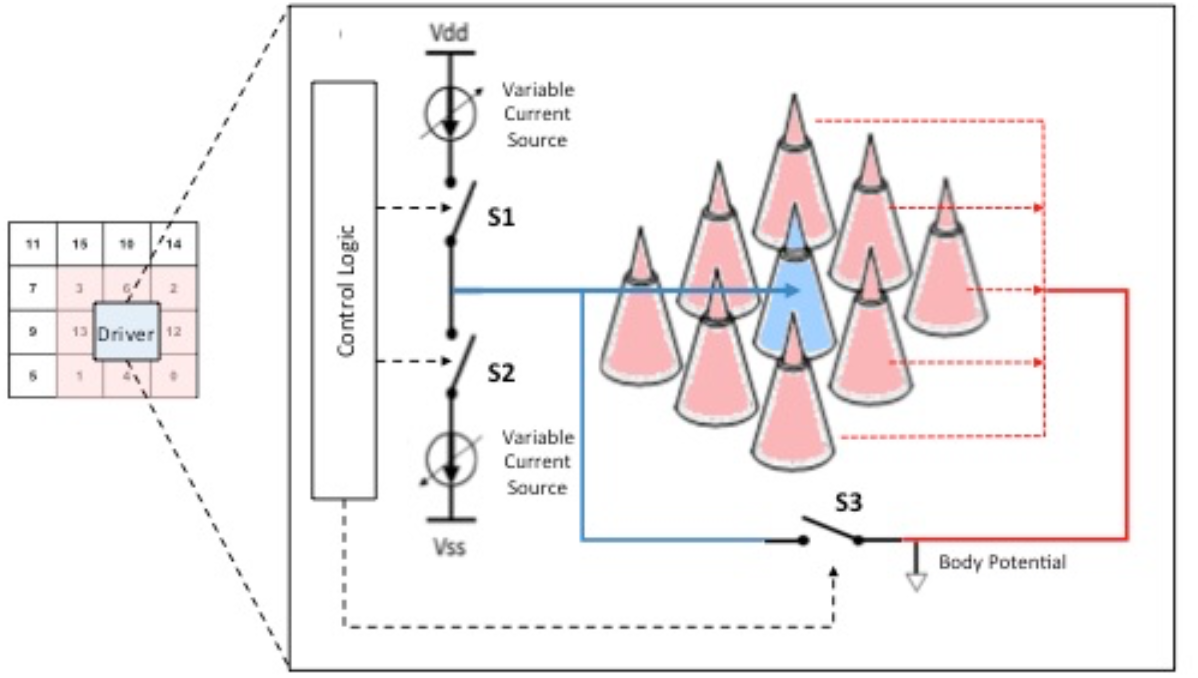
Charge injecting circuitry.

### 2.2.2 Digital Low Pass Filter

The digital low pass filter (DLPF) functions as a second-degree filtration of flickering visual input. It is intended to compensate for quantization errors that originate from the comparator (CMP) filtration of global flickering. There are 576 DLPFs, one per pixel, that accumulate the digitized frame data. Each DLPF is realized as a counter which is increased or decreased for ‘1’ or ‘0’ input, respectively. Typically, it takes seven consecutive frames in which the pixel’s imager data is stable high or low for the pixel’s output to be modified to high (“white”) or low (“black”) accordingly.

### 2.2.3 Receiver and Configuration Manager

The NR600 features configurable settings for both the imager processor and the stimulation logic accounting for specific operation parameters (i.e. pulse duration, amplitude, frequency) which are expected to vary from one patient to another. The NR600 features configurable settings for both the imager and the stimulation logic. These settings can be accessed via a dedicated communication channel, comprised of modulated IR light. The IR modulated illumination is decoded by the receiver and serially transferred to the configuration manager. The communication protocol is Manchester coded, as per IEEE 802.3, with supported rate of 5.6-7.5 Kbps (typically 6 Kbps).

The employed communication protocol comprises of synchronization (min 8 bits), password (8 bits), configuration data (72 bits) and error detection (8 bits CRC). The password serves also as an address; it determines which module is accessed (imager/stimulation logic). Each of the two configurable modules features a 72-bit volatile storage for the configuration parameters. For safety reasons, the default configuration (at startup) prohibits stimulation. A valid configuration transaction is therefore a condition to stimulation.

### 2.2.4 IR Filter

The NR600 IR wavelength used for powering the system is within the imager’s photo-diodes’ sensitivity range. To avoid optical noise, a dedicated IR filter covers the entire imager, with IR attenuation of at least 5 orders of magnitude.

### 2.2.5 Energy

The typical power usage of the device is 300 µW and at extreme stimulation levels it can reach 700 µW. The implant relies on photovoltaic elements embedded in the implant generating operating voltage from an infra-red laser mounted on the NR600 eyeglasses. The wireless and rechargeable eyeglasses provide the implant with power and configuration messages that allow the patient to fine-tune various operating settings.

In the NR600 system, the IR illumination serves both as power and unidirectional communication supply. The Manchester signal duty cycle is 50%, regardless of the transmitted data, therefore the supplied power is in fact limited to half of the peak optical power. The peak intensity (time and space) of the IR illumination is 100 mW/cm^2^, but the receiver remains fully functional, even with IR intensities which are ten times lower.

### 2.2.6 Stimulation pattern

Each electrode is driven by a separate programmable current source. The generated stimuli are symmetric bipolar pulses. Global parameters allow for precise control over the amplitude (0.5-32 µA) and length (50-400 µs) of each pulse, as well as the inter-phase delay (0-350 µs). Local parameters allow further adjustment of the current amplitude for predefined electrodes. Using this method ensures precise control over the transferred charge, while voltage conformance (i.e., water window) is guaranteed by the voltage limited power source (<1 V). The pulses can be combined into stimulation ‘trains’, controlling both the frequency of trains (1-20 Hz) and the number of pulses per train (1-4). Fig. 5 presents a scheme of the stimulation pulse train.

**Figure 5:**
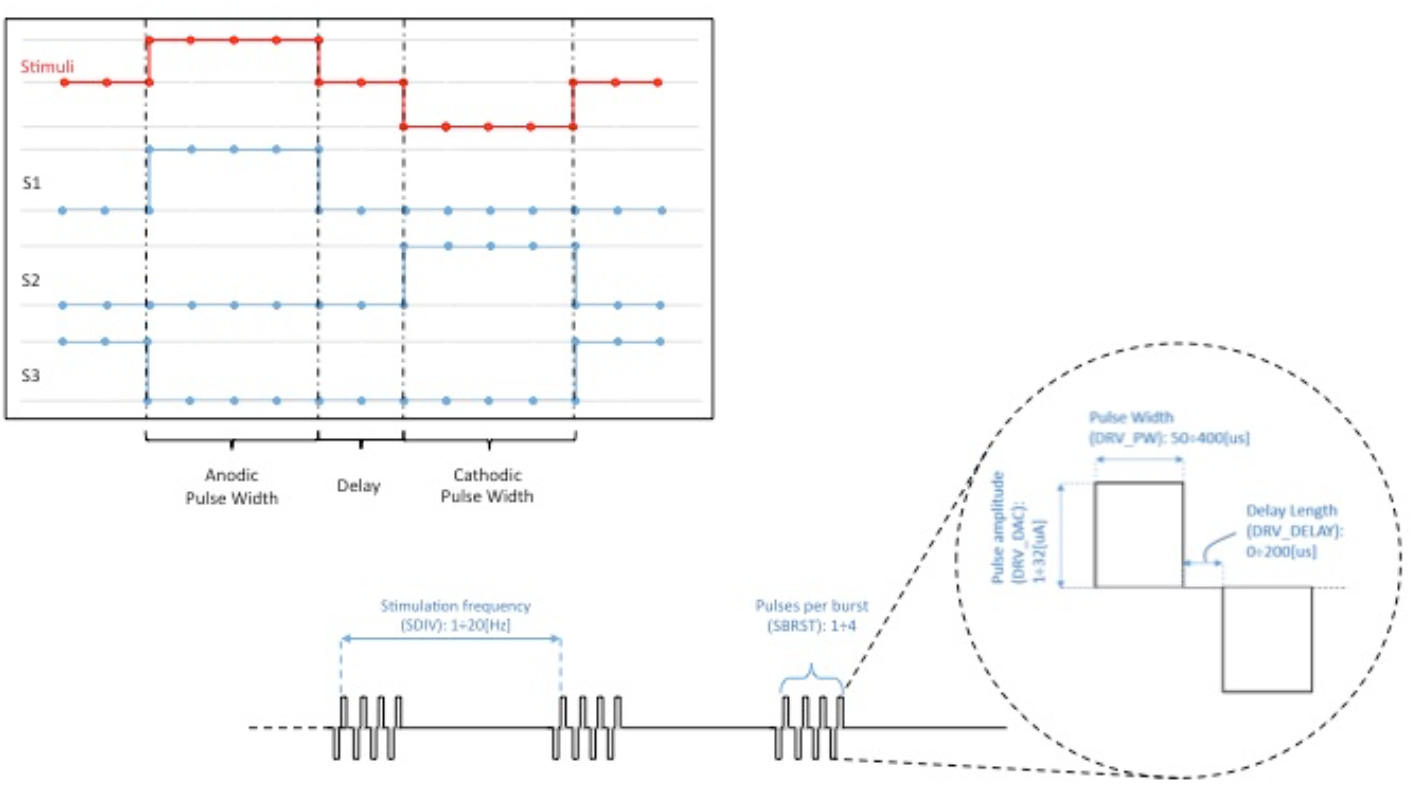
Stimulation pulse parameters can be controlled to optimize stimulation efficacy.

## 3. Surgical Procedure

Owing to its small dimensions, the NR600 implant is inserted into the eye in a minimally invasive surgical procedure lasting approximately two hours. The surgical procedure uses standard ophthalmic surgical techniques of lensectomy and vitrectomy to prepare the eye. To facilitate anchoring to the retina, the NR600 is mounted onto a helical shaped anchoring unit (Fig. 6a). The NR600 implant system is inserted with the helix in its folded state through a limbal incision to the anterior segment of the eye. The incision is sutured and the implant system is anchored to the ciliary sulcus with haptics, resembling intra ocular lens (IOL) fixation. The flexible helical spring is then released and the surgeon guides the implant to its pre-determined site on the macula. As a final step, an IOL is placed on the upper helical ring. The electrodes penetrate the retina with the force of the helix alone, no manual uncontrolled pressure is applied. A key advantage of this anchoring concept is the exclusive contact between the electrode array and the retina, no additional aids such as tacks are needed to hold the implant in place. This reduces the risk of retinal damage that has previously been associated with epiretinal devices, such as retinal tear or detachment [16]. To accommodate the large majority of the patient population, the helix is produced in three different lengths to fit a wide range of eye sizes, 20-26 mm in axial lengths. Since the NR600 implant contains all the necessary functionalities to stimulate retinal nerves, there is no need for additional wiring outside the eye, resulting in a relatively low risk for the procedure and a fast healing and recovery time.

**Figure 6:**
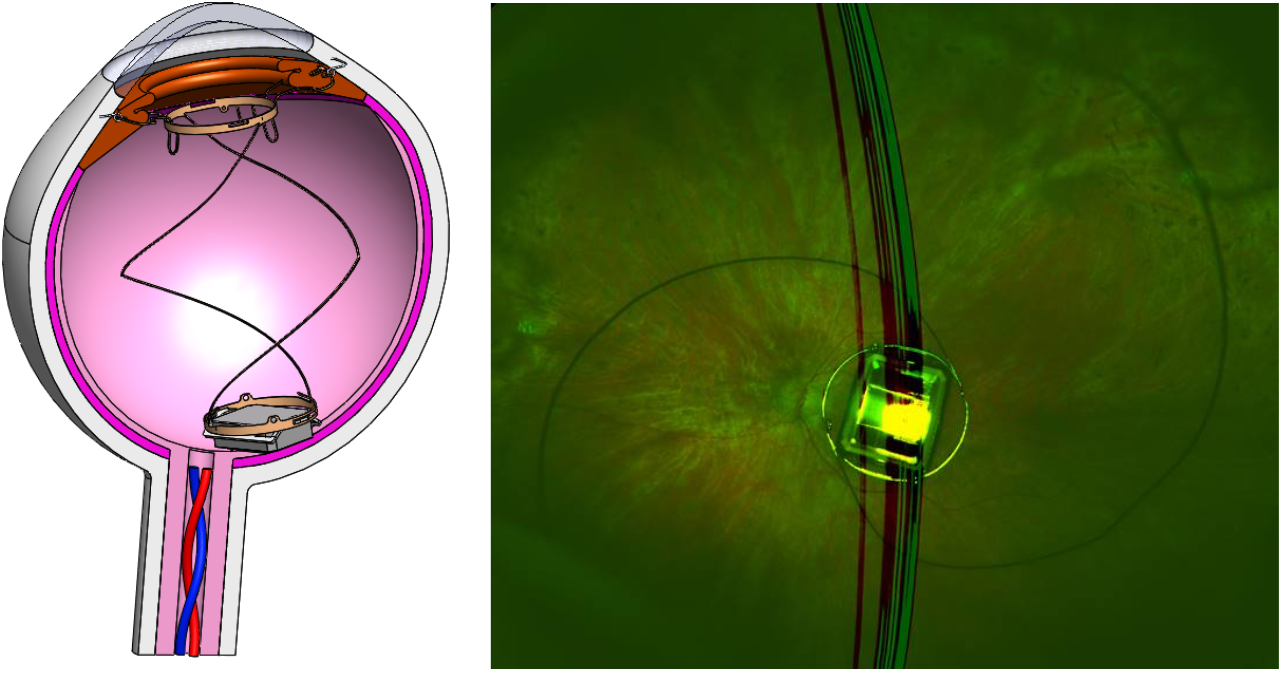
Surgical procedure (a) Implant anchoring scheme consists of a helix shaped holder. (b) Fundus camera image of an implanted device in a human patient, showing the NR600 and the helix anchor.

## 4. First in Human study – Preliminary results

### 4.1 Safety

The NR600 is now being investigated in a feasibility human study. To date (September 2022), nine patients have undergone the implantation procedure. All healed from the surgery remarkably well and were released home after a pre-scheduled brief hospitalization period of 48 hr. Adverse events included one case of mild corneal edema along with slightly elevated intra-ocular pressure (IOP), which commonly occur following ophthalmic surgery. This condition was treated with topical medication and resolved within 3 weeks without a sequel. In another patient there was a case of IOL luxation with subsequent elevated IOP that were resolved by repositioning of the IOL. In follow-ups of up to two years the implanted eyes show no adverse events and are completely clear (Figure 6b).

### 4.2 Visual Perception

All nine patients could perceive phosphenes following activation of the device with relatively low current amplitude and charge injection values (average activation threshold 4.9±2.0 µA and average charge of 1.3±0.7 nC). Visual acuity and visual performance were evaluated starting from one-month post surgery and individual optimization of the device was performed. An example of evaluation results is detailed here. One way to estimate visual acuity is the Visual Grating test, previously performed by both Second Sight and Retina Implant [7], [17]. This is a 4-alternative force choice test in which the patient indicates the orientation (horizontal, vertical, diagonal right or diagonal left) of high contrast lines presented to the subject on a screen 60 cm away. The software presents stripes at different gratings according to an algorithm that increases the spatial frequency in response to correct answers and decreases the spatial frequency in response to wrong answers. Fig. 7 demonstrates a correct answer percentage at different visual gratings on (29±0.8%) versus system off (3±0.4%).

**Figure 7:**
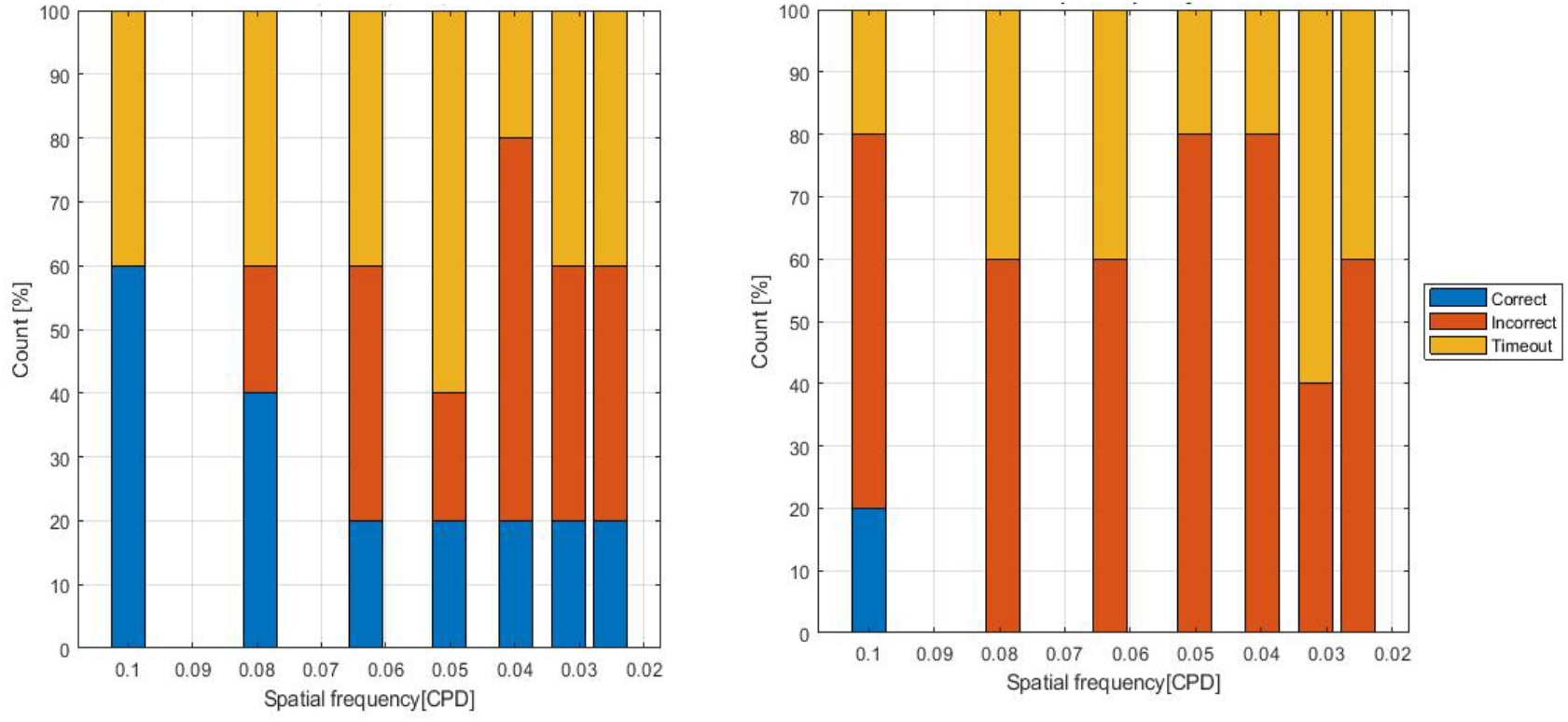
Visual grating visual test result of patient 103, 1-month post-surgery. (Left) NR600 system on. (Right) NR600 off. Blue is for correct, red incorrect and yellow is time out. Data are presented as percentages.

The Square Localization test [7] was implemented to provide an objective measure of spatial vision. Fig. 7a shows square localization results of patient 103. The subject was seated 40 cm away from the center of a 24″ touch screen monitor. In this test, a high contrast, white 1.6” square is presented in random locations on the monitor. When prompted, the subject scans the monitor and attempts to locate the square, touching the screen at the location of the square center. The response error (the distance between the patient’s touch and the center of the target square) is recorded and averaged over forty trials. The results demonstrate a significant improvement of 6.9±10.1 visual degrees at the 1-month follow-up versus 21.4±9.4 at the baseline measurements.

To evaluate how the spatial vision translates to daily visual performance, the ‘walk the line’ test was employed [16]. In this task, the patient is asked to first identify a white line marked on black mats tiled together, which can randomly be in front, or on his/her right or left side. Then, the patient is asked to walk along the line on a 6 m route which is either straight or includes a 90° turn. Figure 8b indicates the path taken by the patient with the NR600 system ON (green) and OFF (red). In this test, the patient used the system to scan the area in front of him/her, locate the line, walk along it, identify the turn and follow the line up to the end of the route. Without the NR600 system, the patient deviated from the line, and it was evident the patient was unable to locate it.

**Figure 8:**
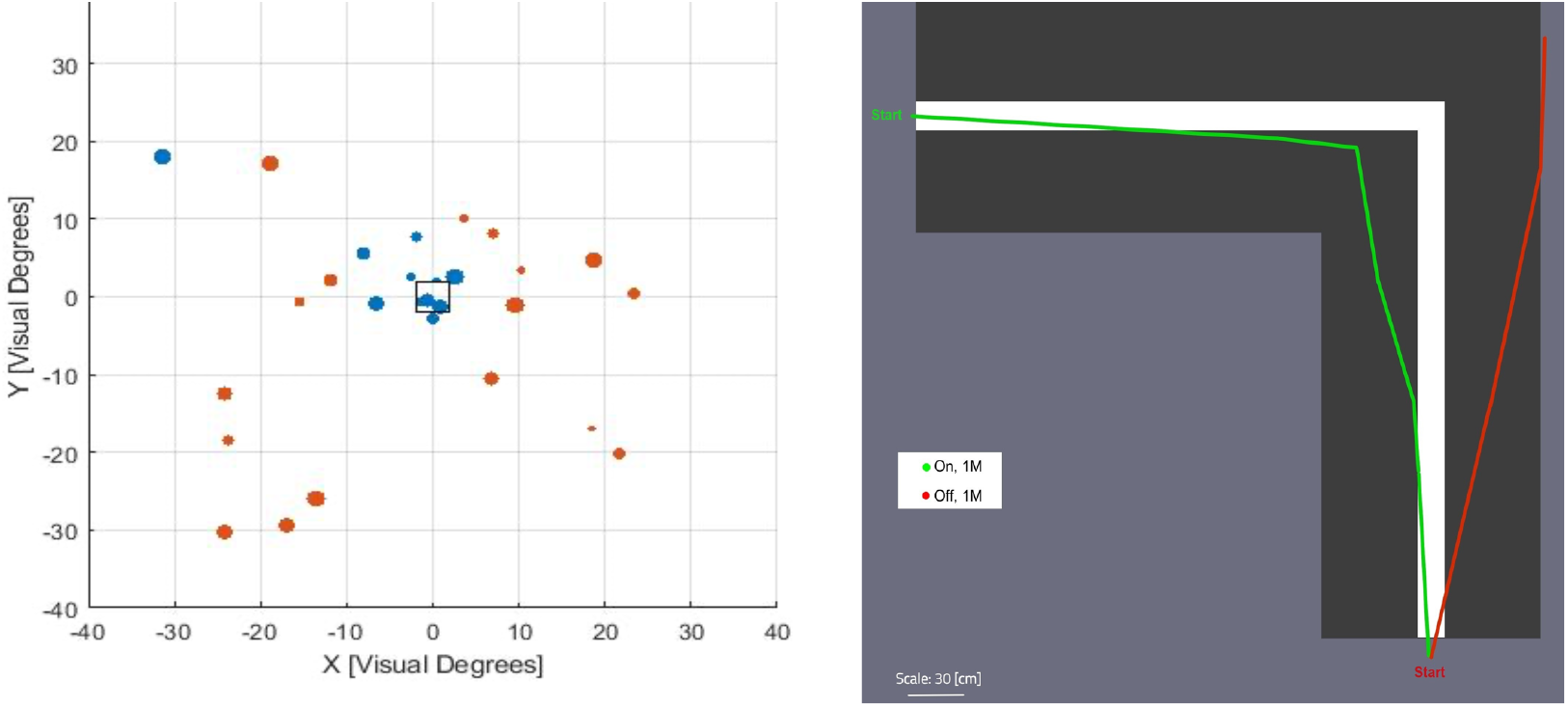
(a) Square localization results. Red dots are at baseline and blue dots are one-month post-surgery when NR600 is ON. The dot size indicates target distance from the screen center. (b) Walk the line task performances of patient 103, one-month post-surgery. Green is with NR600 on and red is off.

An additional test to evaluate functional resolution was the card identification test, in which the patient was asked to identify high contrast patterns of dots, lines and letters. The patient’s success rate was 50% with the NR600 system on versus 8% with the system off. Finally, image detection was tested by ‘locate an object’ test. The patient was asked to identify whether a small object (a key) was placed on a table, as well as locate its position, reach out and pick it up. In this test the patient scored 76% success rate with the NR600 system on versus random guessing with the system off.

## 5. Summary

In this paper we presented the NR600 retinal prosthetic device. The device effectively transduces a visual signal into electrical signals that are delivered to the retina via an array of needle shaped electrodes. We also presented preliminary results from human subjects to demonstrate the ability of the device to generate visual perception in blind subjects.

## 6. Supplementary Videos

Link: http://www.nanoretina.com/

